# Oral health-related quality of life in elderly women participating in a coexistence group in southern Brazil: Oral health and quality of life in elderly women

**DOI:** 10.1101/331066

**Authors:** Thaís Cauduro Dallasta, Vanessa Bischoff Medina, Loiva Beatriz Dallepiane

## Abstract

The objective of this work was to evaluate the association between quality of life with the oral health in elderly women participating in a coexistence group in Southern Brazil. Study of the descriptive type, analytical, cross-sectional approach, with women aged 60 years or more, participants of a coexistence group in a city in Southern Brazil. Data collection used the instrument Oral Health Impact Profile (OHIP-14). The study had the participation of 64 elderly women aged between 60 and 88 years old with a mean of 69.8 ± 7.31 years. The areas that presented the highest values were “Physical Pain”, “Psychological Distress” and “Physical Disability”. The highest averages of the total scores of the OHIP-14 occurred in individuals with lower family income and low education, who showed signs of depression, changes in taste, difficulty to feel the taste of certain foods and malnutrition. Low education, change of taste and malnutrition by the arm circumference were associated with poor quality of life arising from oral disorders.

## Introduction

Oral health problems are recognized as important causes of a negative impact on daily activities, causing pain, suffering, psychological constraints and social isolation. A decreased perception of oral condition can lead to lack of daily oral care, need for dental treatment and, consequently, a poor oral health, affecting quality of life [1].

The Brazilian regions present great inequality in the utilization of dental care and in people’s oral health condition if we consider the access to services. The most vulnerable groups, such as rural populations, elders and poor people, with less schooling, have the worst oral health conditions and face more obstacles in the use of health services [2-3].

The factors associated with the non-utilization of dental services are sex, race/skin color, schooling, income, health insurance, lack of self-perception of oral health and absence of teeth. These factors were also associated with not going to dental consultations for more than 12 months, and other predisposing characteristics, such as age, social networking, and feeding difficulties by oral health problems [4].

Therefore, the social interaction of the elderly and its influence on oral health, and, consequently, on their quality of life, become important. In Brazil, from the concept of active aging, Third Age coexistence groups emerged with places for social gathering as well as occupation of free time with physical and leisure activities among the elderly [5]. These activities provide elderly people integration with a social network with a healthier lifestyle, enabling improvements in health that hindered activities of daily life, thus influencing in a better quality of life [6]. These groups are characterized predominantly by women, following a historic trend in different Brazilian cities, although it is also open to male participation [7].

In this sense, this study aims to assess the association between quality of life with the oral health in elderly women participating in a coexistence group in Southern Brazil.

## Methods

### Study design and sample

The research was characterized as a study of the descriptive type, analytical, with cross-sectional approach, involving female individuals aged 60 years or more, participants of a coexistence group in a city in Southern Brazil.

Data collection occurred in the period between November and December 2015, at the place of activities of the group, being held in the form of an interview. The Research Ethics Committee of the Federal University of Santa Maria - UFSM - approved the research under the opinion number 1,282.020. All participants signed the Informed Consent Form.

### Dependent and independent variables

The dependent variable used in this study was the instrument *Oral Health Impact Profile - OHIP-14* -, which lists the aspects of quality of life most affected by the oral health condition [8].

The questionnaire consists of 14 questions, two for each one of the seven dimensions of oral impact: functional limitation (difficulty in speech and in the decreased sensitivity of taste); pain (sensation of pain and discomfort in the act of eating); psychological distress (concern and stress that the oral condition can cause); physical impairment (possible loss in food and the need to interrupt meals); psychological impairment (difficulty relaxing and feeling of shame on the oral condition); social disability (impact of oral condition in relations with others and the difficulty performing daily activities); and impairment (person’s perception about the impact of the oral condition in his/her life and the inability to develop his/her daily activities).

The questions relate to general oral health problems that people have experienced in the past 12 months, with options for answers on a scale ranging from zero (never) to four (always). As each domain has two subdomains, the score ranges from zero to eight in each domain. The severity is measured by the sum of all scores ranging from zero to 56, since the higher the score the greater is the impact of oral disorders on quality of life.

The independent variables used in this study were sociodemographic variables (family income and schooling) and health conditions (presence of depression, use of dental prosthesis, discomfort while using prosthesis, change in taste, difficulty feeling the food taste and arm circumference).

### Statistical analysis

The data were analyzed using Stata 13.0, presented descriptively for sociodemographic and health characteristics, as well as the mean values of the OHIP-14 scores and their respective domains. The differences between the average scores of the OHIP-14, according to the sociodemographic and health variables, were statistically compared by the Mann-Whitney test, adopting a significance level of 5%. In this study, the scores of the OHIP-14 (outcome) were considered as counting variables, and simple and multiple Poisson regression models were used to verify their association with the other predictor variables. The analysis resulted in the calculation of the means ratio with their respective confidence intervals (RR; IC95%) as an association measure. The RR corresponds to the reason of the arithmetic average of the OHIP-14 between different categories of the predictor variables. The construction of multiple model considered as an entry criterion only the variables that had a p value less than 0.20 in the simple analysis; and they remained in the final model when the p value was less than 0.05 after adjustment.

## Results

Sixty-four elderly women participated in the study. The age ranged from 60 to 88 years, with an average of 69.8 ± 7.31 years. Table 1 presents the sociodemographic characteristics and health conditions of the sample.

**Table 1.** Sociodemographic characteristics and health conditions.

| Variables | N | % |
| --- | --- | --- |
| **Family Income (MW)\*** |  |  |
| ≤ 3 MW | 41 | 64.06 |
| > 3 MW | 23 | 35.94 |
| **Education (years of study)** |  |  |
| ≥ 5 years | 52 | 81.25 |
| < 5 years | 12 | 18.75 |
| **Depression (self-reported)** |  |  |
| No depression | 38 | 80.85 |
| With depression | 9 | 19.15 |
| **Use of prosthesis** |  |  |
| Yes | 55 | 85.94 |
| No | 9 | 14.06 |
| **Discomfort with the prosthesis** |  |  |
| Yes | 30 | 53.57 |
| No | 26 | 46.43 |
| **Change in taste** |  |  |
| No | 49 | 76.56 |
| Yes | 15 | 23.44 |
| **Difficulty feeling food taste** |  |  |
| No | 53 | 84.13 |
| Yes | 10 | 15.87 |
| **Arm circumference** |  |  |
| Depletion | 10 | 15.63 |
| Overweight | 20 | 31.25 |
| Eutrophy | 34 | 53.13 |

\*MW=Minimum wage (Brazilian MW value in November 2015 = R\$ 788,00 or its corresponding at that time, roughly U\$ 203).

Table 2 shows the mean values and the variation in the scores of the OHIP-14 in the sample. The values for the total scores of the OHIP-14 ranged from zero to 28, with an average of 9.78 and standard deviation of 6.66. The scores varied widely according to each domain, and there was no ceiling effect in the answers. The areas that presented the highest values were “Physical pain”, “Psychological distress” and “Physical impairment”.

**Table 2.** Descriptive analysis of the total scores and subdomains of the OHIP-14.

|  | **Number of questions** | **Scores meand (SD)** | **Possible range** | **Observed range** |
| --- | --- | --- | --- | --- |
| OHIP-14 (total score) | 14 | 9.78 (6.66) | 0 – 56 | 0 – 28 |
| **Subdomains** |  |  |  |  |
| Functional limitation | 2 | 1.05 (1.47) | 0 – 8 | 0 – 6 |
| Physical pain | 2 | 1.98 (1.41) | 0 – 8 | 0 – 6 |
| Psychological discomfort | 2 | 1.83 (1.92) | 0 – 8 | 0 – 6 |
| Physical impairment | 2 | 1.63 (1.53) | 0 – 8 | 0 – 5 |
| Psychological impairment | 2 | 1.41 (1.45) | 0 – 8 | 0 – 6 |
| Social impairment | 2 | 1.08 (1.30) | 0 – 8 | 0 – 4 |
| Impairment | 2 | 0.81 (1.33) | 0 – 8 | 0 – 4 |

There was statistically significant difference between the averages of the total scores and subdomains OHIP-14 according to the health conditions variables and sociodemographic characteristics (Table 3). The highest means of the total scores of the OHIP-14 occurred in individuals with lower family income, low educational attainment, with depressive signs, changes in taste, malnutrition and with difficulty to feel the taste of certain foods. Education also associated with OHIP-14 scores in the domains of psychological distress and physical impairment. Similarly, changes in taste also influenced the fields of functional limitation and physical impairment. There were similar results regarding difficulty in feeling the taste of foods, since this variable influenced negatively the domain functional limitation and psychological impairment. Malnourished individuals, according to the arm circumference, also had a worsening of quality of life when compared to individuals with excess weight in the total scores and in the field of physical impairment.

**Table 3.**
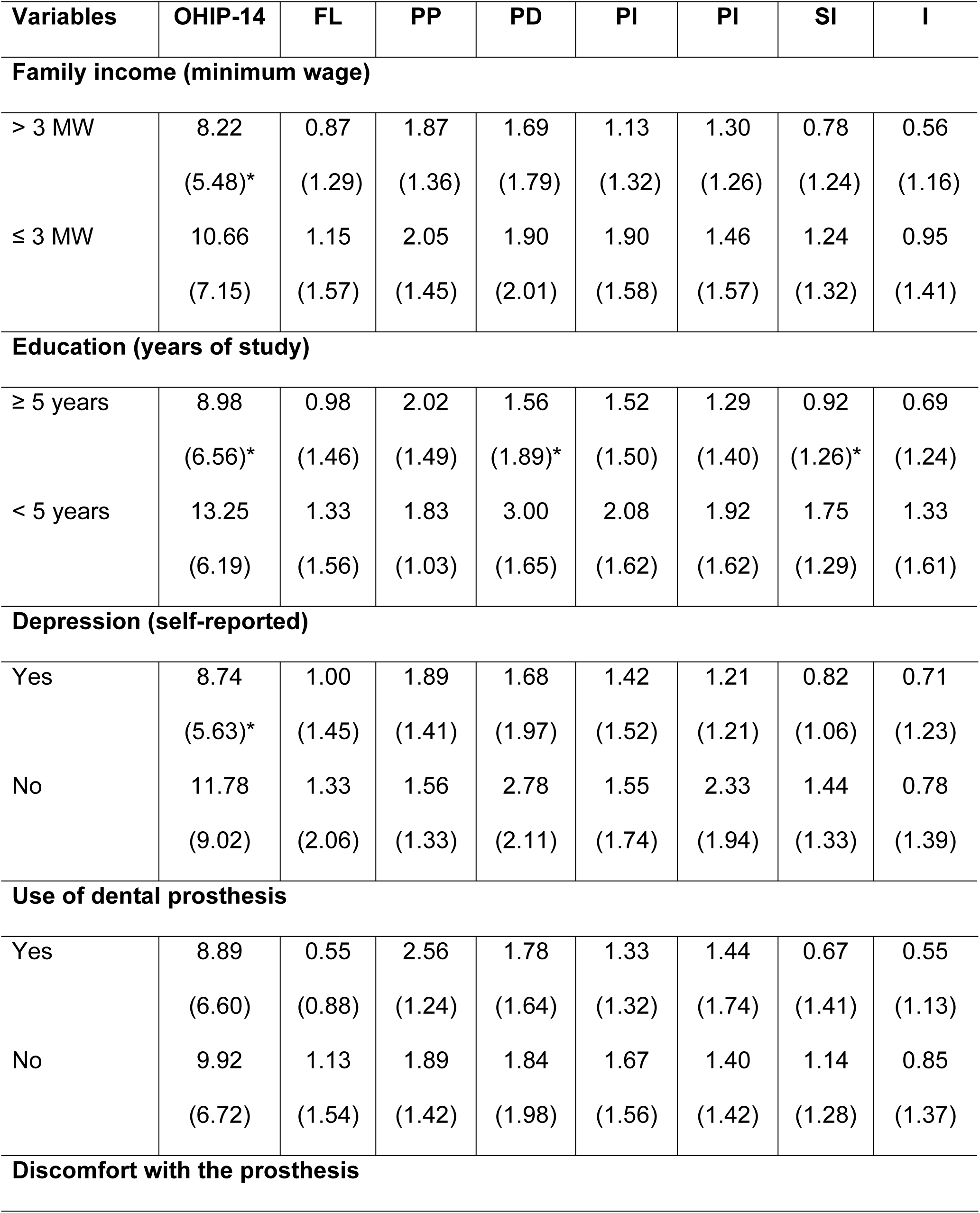
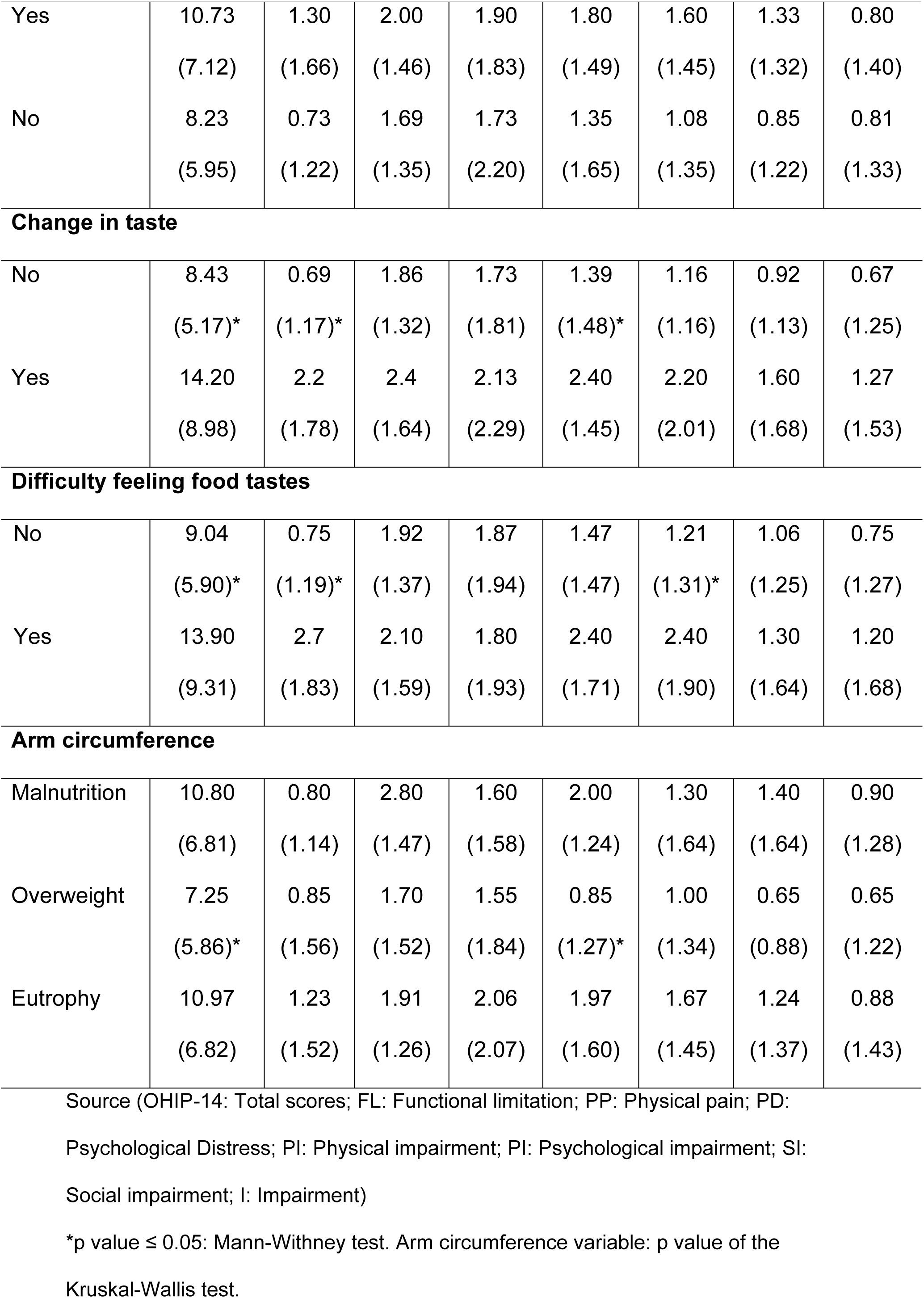
Means (standard deviations) of the total scores and subdomains of the OHIP-14 according to health condition and sociodemographic variables.

| Variables | OHIP-14 | FL | PP | PD | PI | PI | SI | I |
| --- | --- | --- | --- | --- | --- | --- | --- | --- |
| **Family income (minimum wage)** |  |  |  |  |  |  |  |  |
| > 3 MW | 8.22<br>(5.48)* | 0.87<br>(1.29) | 1.87<br>(1.36) | 1.69<br>(1.79) | 1.13<br>(1.32) | 1.30<br>(1.26) | 0.78<br>(1.24) | 0.56<br>(1.16) |
| ≤ 3 MW | 10.66<br>(7.15) | 1.15<br>(1.57) | 2.05<br>(1.45) | 1.90<br>(2.01) | 1.90<br>(1.58) | 1.46<br>(1.57) | 1.24<br>(1.32) | 0.95<br>(1.41) |
| **Education (years of study)** |  |  |  |  |  |  |  |  |
| ≥ 5 years | 8.98<br>(6.56)* | 0.98<br>(1.46) | 2.02<br>(1.49) | 1.56<br>(1.89)* | 1.52<br>(1.50) | 1.29<br>(1.40) | 0.92<br>(1.26)* | 0.69<br>(1.24) |
| < 5 years | 13.25<br>(6.19) | 1.33<br>(1.56) | 1.83<br>(1.03) | 3.00<br>(1.65) | 2.08<br>(1.62) | 1.92<br>(1.62) | 1.75<br>(1.29) | 1.33<br>(1.61) |
| **Depression (self-reported)** |  |  |  |  |  |  |  |  |
| Yes | 8.74<br>(5.63)* | 1.00<br>(1.45) | 1.89<br>(1.41) | 1.68<br>(1.97) | 1.42<br>(1.52) | 1.21<br>(1.21) | 0.82<br>(1.06) | 0.71<br>(1.23) |
| No | 11.78<br>(9.02) | 1.33<br>(2.06) | 1.56<br>(1.33) | 2.78<br>(2.11) | 1.55<br>(1.74) | 2.33<br>(1.94) | 1.44<br>(1.33) | 0.78<br>(1.39) |
| **Use of dental prosthesis** |  |  |  |  |  |  |  |  |
| Yes | 8.89<br>(6.60) | 0.55<br>(0.88) | 2.56<br>(1.24) | 1.78<br>(1.64) | 1.33<br>(1.32) | 1.44<br>(1.74) | 0.67<br>(1.41) | 0.55<br>(1.13) |
| No | 9.92<br>(6.72) | 1.13<br>(1.54) | 1.89<br>(1.42) | 1.84<br>(1.98) | 1.67<br>(1.56) | 1.40<br>(1.42) | 1.14<br>(1.28) | 0.85<br>(1.37) |
| **Discomfort with the prosthesis** |  |  |  |  |  |  |  |  |
| Yes | 10.73<br>(7.12) | 1.30<br>(1.66) | 2.00<br>(1.46) | 1.90<br>(1.83) | 1.80<br>(1.49) | 1.60<br>(1.45) | 1.33<br>(1.32) | 0.80<br>(1.40) |
| No | 8.23<br>(5.95) | 0.73<br>(1.22) | 1.69<br>(1.35) | 1.73<br>(2.20) | 1.35<br>(1.65) | 1.08<br>(1.35) | 0.85<br>(1.22) | 0.81<br>(1.33) |
| Change in taste |  |  |  |  |  |  |  |  |
| No | 8.43<br>(5.17)* | 0.69<br>(1.17)* | 1.86<br>(1.32) | 1.73<br>(1.81) | 1.39<br>(1.48)* | 1.16<br>(1.16) | 0.92<br>(1.13) | 0.67<br>(1.25) |
| Yes | 14.20<br>(8.98) | 2.2<br>(1.78) | 2.4<br>(1.64) | 2.13<br>(2.29) | 2.40<br>(1.45) | 2.20<br>(2.01) | 1.60<br>(1.68) | 1.27<br>(1.53) |
| Difficulty feeling food tastes |  |  |  |  |  |  |  |  |
| No | 9.04<br>(5.90)* | 0.75<br>(1.19)* | 1.92<br>(1.37) | 1.87<br>(1.94) | 1.47<br>(1.47) | 1.21<br>(1.31)* | 1.06<br>(1.25) | 0.75<br>(1.27) |
| Yes | 13.90<br>(9.31) | 2.7<br>(1.83) | 2.10<br>(1.59) | 1.80<br>(1.93) | 2.40<br>(1.71) | 2.40<br>(1.90) | 1.30<br>(1.64) | 1.20<br>(1.68) |
| Arm circumference |  |  |  |  |  |  |  |  |
| Malnutrition | 10.80<br>(6.81) | 0.80<br>(1.14) | 2.80<br>(1.47) | 1.60<br>(1.58) | 2.00<br>(1.24) | 1.30<br>(1.64) | 1.40<br>(1.64) | 0.90<br>(1.28) |
| Overweight | 7.25<br>(5.86)* | 0.85<br>(1.56) | 1.70<br>(1.52) | 1.55<br>(1.84) | 0.85<br>(1.27)* | 1.00<br>(1.34) | 0.65<br>(0.88) | 0.65<br>(1.22) |
| Eutrophy | 10.97<br>(6.82) | 1.23<br>(1.52) | 1.91<br>(1.26) | 2.06<br>(2.07) | 1.97<br>(1.60) | 1.67<br>(1.45) | 1.24<br>(1.37) | 0.88<br>(1.43) |

Source (OHIP-14: Total scores; FL: Functional limitation; PP: Physical pain; PD:
Psychological Distress; PI: Physical impairment; PI: Psychological impairment; SI:
Social impairment; I: Impairment)
\*p value $\leq 0.05$ : Mann-Whitney test. Arm circumference variable: p value of the
Kruskal-Wallis test.

Table 4 presents the results of the simple and adjusted Poisson regression models. The simple analysis showed that only the variables “use of prosthesis and discomfort while using prosthesis” were not associated with the outcome. After adjustment, the variables “educational attainment, change in taste and arm circumference” remained statistically associated to the total score means of the OHIP-14. For example, individuals who had less than five years of study presented a OHIP-14 mean 1.67 times greater when compared to individuals who had five or more years of study (RR: 1.67; 95% IC: 1.29-2.16). In the same way, subjects with change of taste and malnutrition presented worse levels of quality of life related to oral health (i.e. higher OHIP-14 means), when compared to subjects without these changes.

**Table 4.** Association between health condition and sociodemographic variables with OHIP-14 means. Simple and Adjusted Poisson Regression Models.

| Variables | RR<sup>a</sup> (95%CI) | P | RR<sup>b</sup> (95%CI) | p |
| --- | --- | --- | --- | --- |
| **Family income**<br>(MW=minimum wage) |  |  | * |  |
| > 3 MW | 1 |  |  |  |
| ≤ 3 MW | 1.29 (1.09-1.53) | 0.03 |  |  |
| **Education (years of study)** |  |  |  |  |
| ≥ 5 years | 1 |  | 1 |  |
| < 5 years | 1.47 (1.23-1.77) | <0.01 | 1.36 (1.13-1.64) | 0.01 |
| **Depression** |  |  | * |  |
| No depression | 1 |  |  |  |
| With depression | 1.35 (1.08-1.68) | <0.01 |  |  |
| **Use of dental prosthesis** |  |  | **\*\*** |  |
| Yes | 1 |  |  |  |
| No | 1.12 (0.88-1.41) | 0.36 |  |  |
| **Discomfort with the prosthesis** |  |  | **\*** |  |
| Yes | 1 |  |  |  |
| No | 0.77 (0.65-1.01) | 0.06 |  |  |
| **Change in taste** |  |  |  |  |
| No | 1 |  | 1 |  |
| Yes | 1.68 (1.43-1.99) | <0.01 | 1.72 (1.45-2.05) | <0.01 |
| **Difficulty feeling food tastes** |  |  | **\*** |  |
| No | 1 |  |  |  |
| Yes | 1.54 (1.27-1.85) | <0.01 |  |  |
| **Arm circumference** |  |  |  |  |
| Malnutrition |  |  | 1 |  |
| Overweight | 0.67 (0.52-0.87) | <0.01 | 0.92 (0.74-1.14) | 0.41 |
| Eutrophy | 1.01 (0.81-1.26) | 0.90 | 0.62 (0.48-0.80) | <0.01 |

RR<sup>a</sup>: Means ratio of the non-adjusted model; RR<sup>b</sup>: Means ratio of the adjusted
model for all variables.
\* Variables excluded from the final model for losing significance after adjustment.
\*\* Variable use of prosthesis was not included in the multiple model for presenting p>0.20 in the non-adjusted analysis.

## Discussion

This study evaluated the aspects of quality of life most affected by the oral health condition in elderly women participating in a coexistence group in Southern Brazil. The main result of this study is that individuals with less schooling (< 5 years), change in taste and malnutrition, as assessed by the arm circumference, presented the worst levels of quality of life associated with oral health.

Investigating people’s perception and position on their health problems is important, because it allows determining the social, cultural and economic influence in their quality of life. The self-perception in oral health, assessed by means of the Oral Health Impact Profile (OHIP-14) using multilevel analysis, identified that individuals of the female sex, with advanced age, worst scores of quality of life and social support, with bad habits, smokers and residents in low-income places, were more likely to report worse self-perceived oral health [9].

The low schooling compromises the understanding of the concept of oral health as part of overall health. In a study, the prevalence of edentulism, use, need and replacement of dental prosthesis showed a precarious condition of elderly respondents, although reporting a great or good perception of their oral health [10].

Changes in taste influenced the domains of functional limitation and physical impairment. More healthy oral conditions contribute to a better perception of flavor, and may stimulate appetite and, consequently, increase caloric intake; this can help prevent nutritional deficiency in elders and improve the overall health and quality of life of these patients [11].

The changes of taste among elders with physical impairment can be explained by their common use of dental prostheses, which changes the chewing function, decreasing the strength to crush and, with it, hindering the bite of food, considering that natural teeth have no longer the same performance [12]. These factors can lead the elderly person to lose his/her desire to eat, chew (due to early fatigue) and pleasure while eating. Faced with these situations, elders realize that chewing is no longer easy and comfortable, and that there is a need to select the food type or their way to consume it.

According to the findings in this study, the relationship of malnutrition with worse scores in OHIP-14 occurs because malnutrition in elders has an obvious impact on their overall health and quality of life.

In this study, the sense of physical pain, psychological distress and physical impairment were associated with oral disorders and, consequently, to a worse quality of life. In another study, the elders that showed greater severity of OHIP-14 also showed greater impairment of mental domain (depression) and quality of life. Oral health, one of the components of quality of life, refers to the individual’s subjective experience on his/her functional, social and psychological well-being [13].

Changes in the psychological aspect, such as depression, social isolation and loneliness, weaken the elder, causing disinterest for activities of daily life and affecting food consumption, which may motivate a growing disinterest in the face of more consistent healthy foods, which causes, therefore, the installation of inadequate dietary habits, characterized by the intake of foods with a smoother texture and, at the same time, poor in nutrients. This gives the appearance of nutritional deficiencies that impair the functioning of various organs, affecting their health and contributing to a worse quality of life [13].

All elderly patients evaluated in this study had no cognitive deficit, and this may be associated to their participation in coexistence groups characterized by several stimuli, motivated by the coexistence with other elderly people with cognitive demands to develop their capacities. In another study, the participation of the elderly population in coexistence groups has proved to be a good occupational therapeutic resource for health prevention and promotion, as well as a possibility of early cognitive intervention with these subjects [14].

Regarding the aspects related to the individual dimension, all individuals in the study were female, which may represent a limitation to possible inferences or generalizations for the whole population. Women are more linked to the care act (personal and family), thus seeking more health services and reporting more morbidities [1].

Another study found no statistically significant differences between men and women regarding the scores obtained in the OHIP-14, and women showed higher levels, especially in the dimensions of physical impairment and psychological distress. This implies that, if the search was performed including elderly men, the results could be different [15].

During several generations, women played a cultural role of responsibility with family care, and, therefore, are more attentive and concerned with their oral health, while, at the same time, they feel a greater need to consult their doctor regarding any change, thus preventing its progression [16].

Elders’ participation in coexistence groups is important, incorporating a social network, allowing them greater satisfaction with life. The improvements relate to health issues, stating that, before attending the groups, they frequently had headaches that prevented them from performing common activities of daily life [6].

Thus, the activities offered by the groups helped the elders in this study to acquire a healthier lifestyle and, consequently, improve their quality of life.

Among the limitations of this research, its cross-sectional design stands out, not allowing establishing cause and effect relationships between associated factors. Nevertheless, the presented results are valid and representative of the investigated elderly population, allowing more clarification about the quality of life related to elders’ oral health. The assessment of quality of life is a dynamic process, and possible associated factors can modify throughout time. Another limitation of the study was the non-completion of clinical assessment to verify the oral situation of the elderly interviewees. Such limitations are some starting points for future researches to fill these gaps.

## Conclusion

Low schooling, change of taste and malnutrition by the arm circumference were associated with poor quality of life arising from oral disorders.

## Acknowledgements

We would like to thank the Postgraduate Program in Gerontology of the UFSM and to the elderly members of the coexistence group “*Alegria de Viver*” by their receptiveness and availability to participate in our research.

